# Basal expression of immune receptor genes requires low levels of the phytohormone salicylic acid

**DOI:** 10.1101/2023.07.14.548351

**Authors:** Tijmen van Butselaar, Savani Silva, Dmitry Lapin, Iñigo Bañales, Sebastian Tonn, Chris van Schie, Guido Van den Ackerveken

## Abstract

The hormone salicylic acid (SA) plays a crucial role in plant immunity by activating responses that arrest pathogen ingress. Since SA accumulation also penalizes growth, the question remains why healthy plants synthesize this hormone. By overexpressing SA-inactivating hydroxylases in *Arabidopsis thaliana*, we reveal that basal SA levels in unchallenged plants are needed for expression of selected immune receptor and signaling genes, thereby enabling early pathogen detection and activation of immunity.

## Main text

Plants activate their immune response to biotrophic pathogens largely through the phytohormone salicylic acid (SA)^1,2^. This encompasses not only the transcriptional activation of defense genes but also the repression of growth and development-related genes^3^ which translates into reduced plant growth^4,5^. This balance or so-called growth-immunity tradeoff must be well-controlled to circumvent complete immunity-driven growth arrest^4,5^. Therefore, plant SA responsiveness should be tightly regulated through SA synthesis, catabolism, and signaling^2^. In *Arabidopsis thaliana* (Arabidopsis hereafter), SA catabolism is largely driven by the Fe(II) oxoglutarate-dependent oxygenases DMR6/S5H (DOWNY MILDEW RESISTANT 6/SA 5-HYDROXYLASE) and its functionally redundant paralog DLO1/S3H (DMR6-LIKE OXYGENASE 1/SA 3-HYDROXYLASE)^6^. DMR6 and DLO1 hydroxylate SA to form 2,5-dihydroxybenzoic acid (2,5-DHBA) and 2,3-DHBA, respectively, which are rapidly glucosylated and transported into plant vacuoles^7-9^. Reduced SA catabolism, as in *dmr6* and *dlo1* single and double mutants, leads to increased SA levels, elevated expression of immunity-related genes, and enhanced broad-spectrum resistance to biotrophic pathogens^6-8^. However, severe growth reduction is observed in the *dmr6 dlo1* double mutant, caused by hyperaccumulation of SA^6^. Conversely, plants overexpressing *DMR6* or *DLO1* have lower SA levels, higher pathogen susceptibility, and increased growth^6-8^. *dmr6*-based disease resistance is not only effective in Arabidopsis but also different crops^10-17^, demonstrating its potential for broad-spectrum resistance breeding.

Perturbation of *DMR6* and *DLO1* in Arabidopsis allows controlling basal SA levels in mutants and overexpression lines and studying effects of SA on growth, immunity, gene expression, and other responses without exogenous application of the hormone^6,18^. Here, a comparison of transcriptomes was carried out on Arabidopsis lines with perturbed expression of *DMR6* and *DLO1* (**Supplementary Figure 1**). Overexpression of *DMR6* or *DLO1* in Arabidopsis Col-0 (hereafter Col) was associated with increased rosette size of the 4.5-week-old plants and lower total SA levels in both 2- and 4.5-week-old plants, in agreement with previous results^6,8^ (**Figure 1A-C)**. Interestingly, the overexpression lines were larger and had lower SA levels at two weeks than the *sid2-1* mutant which has strongly reduced SA production (**Figure 1A-C**). Single *dmr6-3* (hereafter *dmr6*) and *dlo1* mutants grew smaller and had increased SA levels compared to Col, whereas the *dmr6 dlo1* double mutant was severely reduced in growth and accumulated high SA levels (**Figure 1A-C**). Due to severe leaf senescence^6,8^, the 4.5-week-old *dmr6 dlo1* mutant was excluded from the RNAseq analysis.

**Figure 1.**
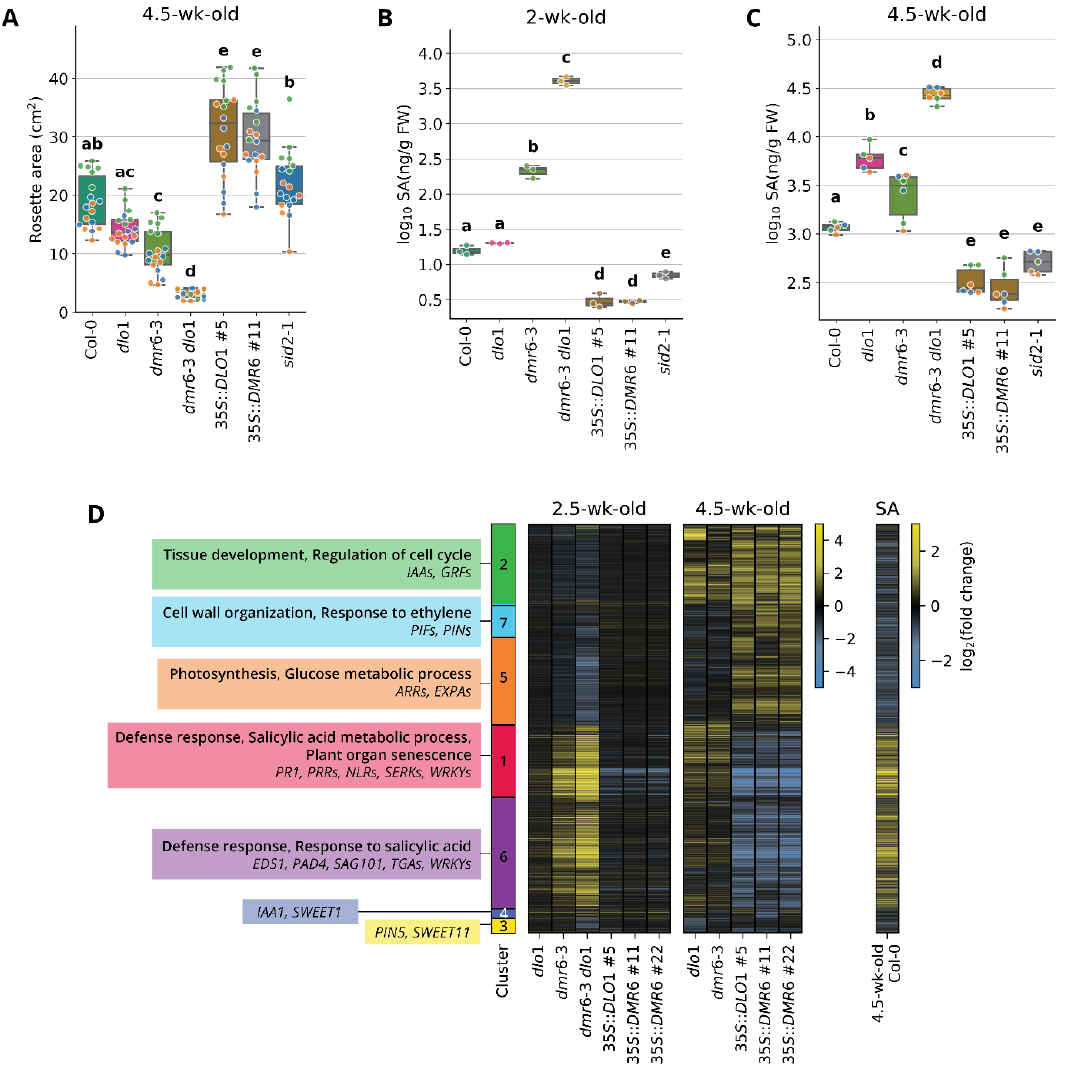
*DMR6/DLO1* perturbation influences Arabidopsis growth, SA levels, and transcriptomes. **(A)** Rosette area of 4.5-week-old plants with mutated or overexpressed *DMR6* and *DLO1* alongside controls Col-0 and *sid2-1*. (**B, C)** Total SA levels in two (**B**) and 4.5 (**C**) week-old plants of indicated genotypes. Data in **A-C** are from three independent experiments (indicated by differently colored dots in **A** and **C;** in **B**, each dot represents an independent experiment). Different letters above the boxplots show statistically significant differences between genotypes (two-way ANOVA (**A, C**) or one-way ANOVA (**B**) followed by Tukey’s post-hoc test, *p* ≤ 0.05). Outliers were removed with interquartile range method (IQR) in **C. (D)** Heatmap of log_2_ fold change values for the 6234 genes that were differentially expressed (|log_2_FC| ≥ 1, FDR-adj. *p* ≤ 0.05) in *DMR6* and *DLO1* mutant or overexpression lines compared to the age-matched wild-type. Selected GO terms enriched in seven clusters of co-expressed genes are listed with example genes or gene families from each cluster. On the right is the RNAseq profile of Col leaves 24 hours after treatment with 0.5 mM SA^20^.

The largest transcriptome changes were detected in the 2.5-week-old *dmr6 dlo1* double mutant and 4.5-week-old *DMR6/DLO1* overexpression lines compared to Col (**Supplementary Figure 1A**). In total, we found 6234 differentially expressed genes (DEG) between the Col control and mutants or overexpression lines of the same age (|log_2_FC|≥1, FDR-adj. *p*≤0.05; **Figure 1D, Supplementary Figure 1B-C**). Hierarchical clustering grouped these genes into seven clusters (**Figure 1D**). Clusters 1 and 6 (1103 and 1705 DEG, respectively) contained genes upregulated in the 2.5-week-old *dmr6* single and *dmr6 dlo1* double mutants but downregulated in 4.5-week-old *DMR6* or *DLO1* overexpression lines. These clusters were enriched for SA and other immunity-related gene ontology (GO) terms (**Figure 1D, Supplementary Figure 2**). In contrast, genes in clusters 2 (1217 DEG), 5 (1328), and 7 (504) were downregulated in the 2.5-week-old *dmr6 dlo1* double mutant and upregulated in 4.5-week-old *DMR6* or *DLO1* overexpression lines. They were enriched for GO terms associated with photosynthesis, growth, and development (**Figure 1D, Supplementary Figure 2**). Similar to these GO enrichment patterns, genes involved in immunity were also upregulated in tomato *dmr6* mutants^12^ and the Arabidopsis *dmr6-1* mutant^19^, while genes involved in photosynthesis, growth, and development were downregulated in tomato *dmr6* mutants^12^.

The highest number of DEG compared to Col was observed in 2.5-week-old *dmr6 dlo1* plants (3696 DEG) (**Figure 1B**). A stronger effect on the transcriptome was observed in young plants of the *dmr6* single mutant than in older plants (666 and 83 DEG, respectively), whereas the opposite pattern was observed in the *dlo1* mutant (8 and 851 DEG), suggesting that DMR6 and DLO1 are relatively more important for SA catabolism in younger and older plants, respectively. The latter observation supports the role of DLO1 as a regulator of senescence induction^8^. This temporal effect of single mutants on the transcriptome is also reflected in SA levels that are higher in *dmr6* than *dlo1* at 2 weeks but higher in *dlo1* than *dmr6* at 4.5 weeks (**Figure 1B-C**)^8^. In the *DMR6* and *DLO1* overexpression lines, SA levels were significantly lower in both 2- and 4.5-week-old plants compared to Col (**Figure 1B-C**). The overexpression lines had minor transcriptome changes relative to Col in 2.5-week-old plants, while they differed significantly at 4.5 weeks (2617 and 2774 in *35S:DMR6*, depending on the line, and 2895 DEG in *35S:DLO1*; **Supplementary Figure 1B-C**). Together, these data suggest that reduced SA levels have an increasing effect on the transcriptome as the plants age. Although DMR6 and DLO1 produce different DHBAs, the transcriptome changes of 4.5-week-old *DMR6* and *DLO1* overexpression lines compared to Col were strongly correlated (R^2^=0.84 to 0.91, *p*-value < 0.001) (**Supplementary Figure 3A**) indicating that it is the removal of SA that drives transcriptome changes in these lines rather than the production of specific DHBAs. To independently validate that transcriptome changes in the tested genotypes are due to perturbed basal SA levels, we analyzed a published gene expression profile of SA-treated Arabidopsis leaves^20^. It correlated positively with the transcriptome changes of the *dmr6 dlo1* mutant (R^2^=0.67, *p*-value < 0.001) and negatively with that of the *DMR6/DLO1* overexpression lines (R^2^=-0.55 to -0.64, *p*-value < 0.001; **Figure 1D, Supplementary Figure 3B**).

When inspecting the DEG clusters (**Figure 1D**) for known regulators of Arabidopsis immunity, we noticed that 4.5-week-old *DMR6/DLO1* overexpression lines had reduced expression of genes involved in pattern-triggered immunity (PTI) and effector-triggered immunity (ETI). Examples are *EDS1* (*ENHANCED DISEASE SUSCEPTIBILITY 1*), its paralogous signaling partners *PAD4* (*PHYTOALEXIN DEFICIENT 4*) and *SAG101* (*SENESCENCE-ASSOCIATED GENE 101*), genes encoding helper nucleotide-binding leucine-rich repeat proteins (*NLRs*), selected immune receptors from the RLP class *(RECEPTOR-LIKE PROTEINs*, including *RLP23*), and RLP co-receptor *SOBIR1* (*SUPPRESSOR OF BIR1 1*; **Figure 2A**). On the other hand, expression of selected pattern recognition receptor genes from the RLK (*RECEPTOR-LIKE KINASE*) class (e.g., *FLS2* (*FLAGELLIN-SENSITIVE 2*)) was not consistently suppressed in the *DMR6/DLO1* overexpression lines (**Figure 2A**).

**Figure 2.**
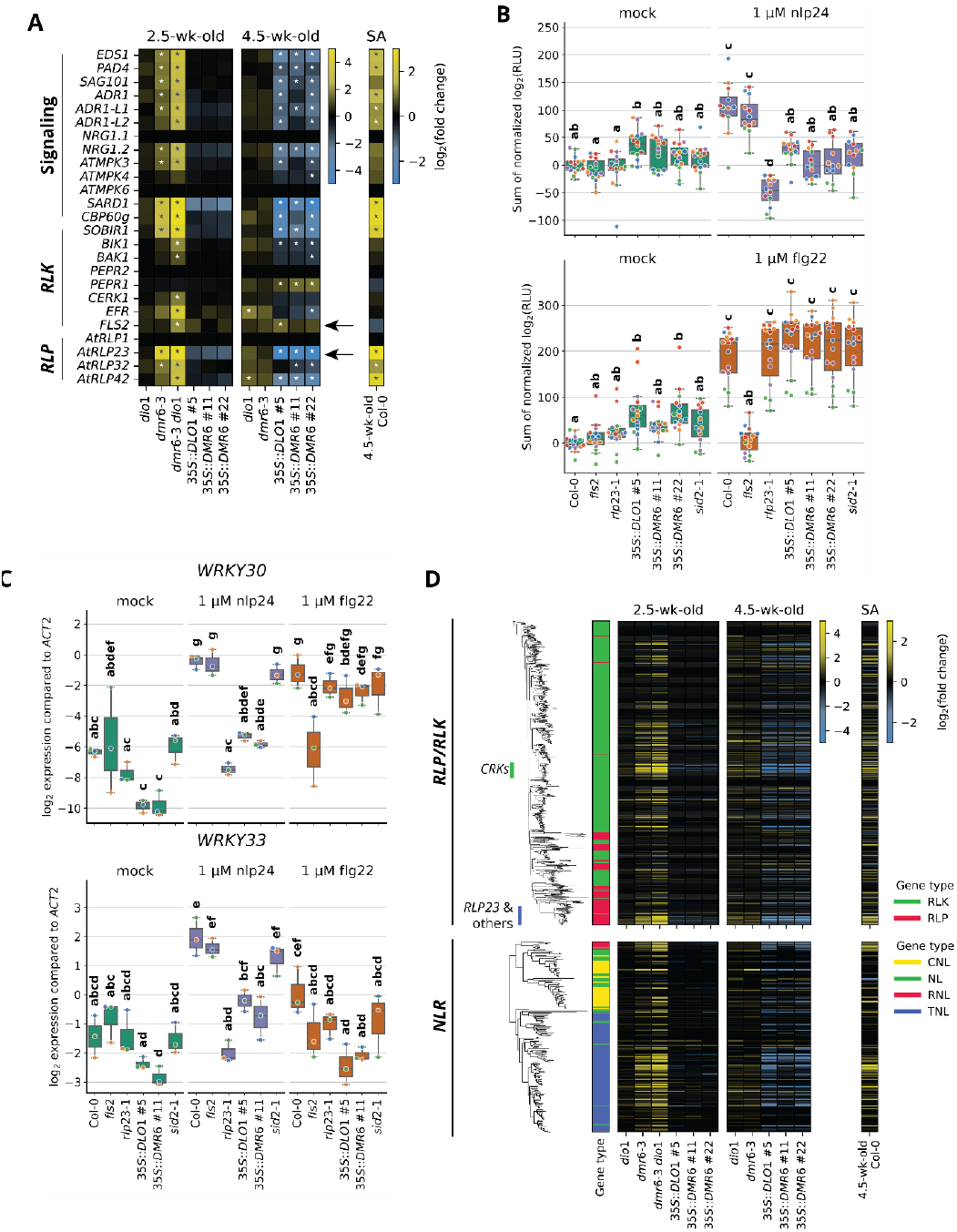
Reduction in the basal SA levels lowers the expression of selected immunity genes and associated PTI responses. **(A)** Expression of PTI-related genes, including signaling components and immune receptors in indicated genotypes. Asterisks denote the differential expression compared to the age-matched Col (|log_2_FC| ≥ 1, FDR-adj. *p* ≤ 0.05). **(B)** Reduced ROS burst in response to nlp24 (upper panel) but not to flg22 (lower) in the SA-depleted lines compared to Col-0. The *rlp23* and *fls2* mutants were negative controls for the nlp24 and flg22 treatments, respectively. RLU: relative luminescence units; mock: mQ. **(C)** Transcript levels of *WRKY30* and *WRKY33* in response to the nlp24 and flg22 treatments. The *DMR6/DLO1* overexpression lines have *WRKY* gene induction to nlp24. *WRKY* transcript levels were measured by qRT-PCR and receptor mutant lines *rlp23* and *fls2* were included as negative controls. Plants in **B** and **C** were 4.5-week-old. Data displayed are derived from three independent experiments, as indicated by differently colored dots. Different letters denote statistically different groups from two-way ANOVA followed by Tukey’s Post—Hoc test, *p* ≤ 0.05. **(D)** Expression of 861 *RLPs* and *RLKs* (top), and 166 *NLRs* (bottom) in *DMR6/DLO1* mutant and overexpression lines. Phylogenetic analysis was performed on protein alignments. Abbreviations: CRK – cysteine-rich receptor-like kinase, CNL, RNL and TNL – NLRs with coiled-coil, RPW8 (RESISTANCE TO POWDERY MILDEW 8), and TIR (Toll/interleukin-1 receptor homology) domains, respectively.

The altered expression of PTI-related genes prompted us to investigate the attenuation of early PTI responses to the pathogen-associated molecular patterns (PAMPs) nlp24 and flg22 in the *DMR6/DLO1* overexpression lines (**Figure 2B-C**). The nlp24-triggered ROS burst and the induction of *WRKY30* and *WRKY33* were strongly reduced in *DMR6*/*DLO1* overexpression lines (**Figure 2B-C**). This fits the lower expression of genes required for the nlp24-induced responses (*RLP23, SOBIR1, EDS1/PAD4/SAG101*, and helper *NLRs*; **Figure 2A**)). The flg22-triggered ROS burst and *WRKY30* and *WRKY33* induction were unaffected in the *DMR6*/*DLO1* overexpression lines, aligning with the unaltered expression of RLKs *FLS2* and *BAK1* (*BRI1-ASSOCIATED RECEPTOR KINASE 1*; **Figure 2A-C**). The single and double *dmr6* and *dlo1* mutants behaved like Col in the tested outputs, suggesting that increasing SA levels does not enhance early PTI responses (**Supplementary Figure 4**). To obtain additional evidence for the role of basal SA levels and SA perception on early PTI responses, we tested the SA-deficient *sid2* mutant (**Figure 1B-C**) and the SA-insensitive *npr1-1 npr4-4D* double mutant^21^. Both mutants showed reduced nlp24-triggered ROS burst (**Figure 2B, Supplementary Figure 5**) confirming the dependency on SA for nlp24 responsiveness.

To obtain a genome-wide view of the effects of *DMR6/DLO1* perturbation on immune receptor gene expression, we analyzed the expression of Arabidopsis genes encoding for RLKs, RLPs, and NLRs in *DMR6*/*DLO1* overexpression lines and mutants. We termed these genes SA-responsive if, compared to the Col control, they were downregulated in the three *DMR6/DLO1* overexpression lines at 4.5 weeks and upregulated in the *dmr6-3 dlo1* double mutant at 2.5 weeks (|log_2_FC|≥1, FDR-adj. *p*≤0.05). We found that 120 *RLPs/RLKs* (14%) and 21 *NLRs* (13%) fell into this group. Notably, the phylogenetic clustering of receptors did not separate SA-responsive genes from the rest (**Figure 2D**), indicating that the dependency of their expression level on basal SA is not restricted to specific phylogenetic clades. We further tested if SA-responsive receptor genes have enrichment of certain transcription factor binding sites. Indeed, promoters of SA-responsive *RLP*/*RLK* and *NLR* genes showed specific enrichment of WRKY transcription factor binding sites (**Supplementary Figure 6**), suggesting that WRKYs contribute to SA dependence of the receptor gene expression.

So far, the role of SA in plant immunity was focused on defense signaling and senescence. Here, we show that low basal levels of SA are important for the appropriate expression of genes encoding for several groups of immunity-related RLPs and their immediate downstream signaling components. Although plants grow better in the absence of basal SA, our results reveal a trade-off in pathogen detection. This explains why low basal SA levels are needed to have a well-functioning plant immune system.

## Data availability

Raw read RNA-seq data have been deposited in the European Nucleotide Archive (ENA) database at EMBL-EBI (https://www.ebi.ac.uk/ena/) under accession number PRJEB61019.

## Acknowledgements

We would like to thank Nora Ludwig (deceased), Joyce Elberse, and Rob Schuurink for the measurements of the hormone data in Figure 1B, part of which was published earlier in Zeilmaker, et al. _6_. This project was funded by Enza Zaden B.V. and Topsector Tuinbouw & Uitgangsmaterialen.

## Author contributions

TvB and SS performed the RNAseq experiment; TvB, SS, DL, and GvdA analyzed RNAseq data; IB conducted the phylogenetic analysis; TvB and CvS measured salicylic acid levels; TvB and DL performed gene expression and ROS burst assays. TvB, DL, and GvdA wrote the manuscript with contributions from all authors.

Authors declare no competing interests.

## Material and Methods

### Plant genotypes and growth conditions

Lines of *Arabidopsis thaliana* (L.) Heynh. Col-0 *dmr6-3, dlo1, dmr6-3 dlo1, 35S:DMR6* and *35S:DLO1* were previously described ^6^. The Col-0 *npr1-1* and *npr1-1 npr4-4D* mutants (Ding, et al. ^21^) were received from Pingtao Ding (Leiden University, the Netherlands). Seeds were imbibed for four days at 4°C, either in 0.1% agarose and then transferred to soil (5:12 sand:soil mix, supplemented with half-strength Hoagland solution, see Van Wees, et al. ^22^) or sown directly on soil. Plants were grown under short-day (10h/14h light/dark, 21°C, 70% relative humidity, 100 μmol/m^2^/sec) conditions with regular watering and a supplement of half-strength Hoagland solution once a week. During the first week of growth, seeds were germinated under 100% RH.

### RNA sequencing library preparation

Aerial parts of 2.5-week-old or the sixth leaf of 4.5-week-old plants were harvested and snap-frozen in liquid nitrogen. RNA isolations and library preparations were performed according to Bjornson, et al. ^23^ high-throughput RNA isolation and library prep protocols, with reagents from other suppliers that are detailed below. Briefly, mRNA was enriched from crude cell lysate in two rounds using biotinylated oligo-dT (IDT Europe) and streptavidin beads (New England Biolabs). RNA isolations were performed in batches in a randomized order. Following cDNA synthesis and adapter ligation, sequencing libraries were generated using indexed primers and enrichment primers (IDT Europe, see Primer List) with Phusion HF polymerase (Thermo Fisher). Libraries were visually inspected on an agarose gel before a double clean up with Ampure XP magnetic beads (Beckman). Library DNA quantity was measured with SYBR Green (Thermo Fisher). Equalized amounts of libraries were pooled together before a final Ampure XP cleanup. Libraries were sequenced at Useq (Utrecht, The Netherlands) on the Illumina NextSeq2000 (P3 1×50nt) platform.

### Transcriptome analysis

All sequencing data analyses on read files were performed in *slurm* workload manager on a local High-Performance Computing Facility (University Medical Center Utrecht, The Netherlands). Illumina BCL files were demultiplexed and converted to fastq format with *bcl2fastq* v2.20.0 (Illumina). Quality of sequencing data was verified before and after trimming with *fastQC* v0.11.5 and *MultiQC* v1.5 ^24^. Trimming was performed with *trimmomatic* v0.39 ^25^ using default settings and Truseq3-PE adapters. Trimmed reads were pseudo-aligned to the TAIRv10 nuclear transcriptome with *kallisto* v0.46.1 ^26^ using a 21-mer index file. Only samples with least 5 million transcript-assigned reads were considered in further analyses. For these samples, the reads per transcript were pooled per gene with *tximport* v1.24.0 ^27^, and differential gene expression was performed with *deseq2* v1.36.0^28^, using a DEG cutoff at |log_2_FC| ≥ 1, FDR-adj. *p* ≤ 0.05.

Fold-change values were further processed in Python v3 using *pandas* 1.4.1 package for data handling and *seaborn* 0.11.2 and *matplotlib* 3.5.1 packages for data visualization. Principal component analysis (PCA) was performed with *scikit-learn* 0.24.2. UpSet plots ^29^ were generated with *upsetplot* 0.61 package for Python using minimal subset size of 25 and minimal degrees of 2. Clustering of genes was performed with hierarchical clustering from *SciPy* 1.7.1 *clusters*.*hierarchy* package, using *complete* method and *correlation* metrics. Clusters were generated with *SciPy* 1.7.1 *hierarchy*.*cut_tree* method, and the optimal *n* clusters for the dataset was determined after visual examination of the plots. Enrichment of biological process gene ontology (GO) terms was performed in ClueGO v2.5.8 plugin for CytoScape^30^. From the enrichment analysis we processed only the overview GO terms to minimize redundancy. Pearson’s correlation analyses were performed with *SciPy* 1.7.1. Enrichment of transcription factor binding sites was conducted in *AME* from the *MEME* 5.4.1-suite ^31^ on gene promoter sequences (1kb upstream of translation start site) (TAIRv10). *AME* search was performed with motifs from DAP-seq database ^32^ and PBM database ^33^ and with an *E*-value threshold of 0.001.

### Phylogenetic analyses

To find groups of sequence-related NLR proteins, the proteome of *A. thaliana* Col-0 (Araport11, representative peptide sequences) was scanned against Pfam-A database (release 35.0, pfam_scan.pl -e_seq 0.1 -e_dom 0.1), and NB-ARC domains (PF00931.25) of the corresponding 166 proteins were extracted for phylogenetic analysis (Biopython v1.79). 92% of manually identified NLRs proteins were supported by Araport11 annotation. Multiple sequence alignments (MSAs) were constructed with MAFFT default parameters (v7.505, --auto)^34^ and filtered for parsimony-informative sites. Alignment columns with >60 % gaps were also removed (Clipkit v1.3.0, -m kpic-gappy -g 0.6). Filtered alignment was inspected in the Wasabi MSA browser (http://was.bi/) prior to phylogeny reconstruction. The ML trees were inferred with IQ-TREE (v.2.1.2, -m MFP -B 1000 -alrt 1000 -T auto)^35^. The best-fit substitution model for the data was determined by ModelFinder (JTT+F+R6)^36^.

For the sequence-based grouping of Arabidopsis RLKs and RLPs, we used 695 RLKs and 175 RLPs annotated in previous publications^37-39^ and from the RGAugury pipeline^40^. Four sequences (AT1G16140, AT4G21370, AT5G57670 and AT3G21960) were discarded since they were annotated as pseudogenes in TAIR10.1. MSAs were constructed with MAFFT (v7.505, --auto) and filtered for parsimony-informative sites. Alignment columns with >90 % gaps were also removed (Clipkit v1.3.0, -m kpic-gappy -g 0.6). Filtered alignment was inspected in the Wasabi MSA browser (http://was.bi/) and the conserved region was extracted for evolutionary inference (seqkit v2.3.0, region between 700:1379 columns). AT5G49750 was removed from further analysis due to the lack of aligned positions in this area. The ML trees were inferred with IQ-TREE (v.2.1.2, -m MFP -B 1000 -alrt 1000 -T auto). The best-fit substitution model for the data was determined by ModelFinder (LG+F+R8). Resulting NB-ARC and RLP/RLK trees were inspected and rooted in iTOL (v6)^41^.

### Salicylic acid measurements

Total SA levels in two-week-old plants are from the experiment published in Zeilmaker, et al. ^6^ where data for the *DMR6* and *DLO1* overexpression lines were unpublished. Total SA quantification on 4.5-week-old plants was performed as follows. Aerial parts of plants were weighed to approximately 200 mg material, harvested in liquid nitrogen, and subsequently freeze-dried overnight. Samples were homogenized with steel beads before extraction with 1 ml 80% ethanol 0.5% formic acid and 3 ppm 5-fluorosalicylic acid (internal standard). Cell debris was spun down and supernatant was evaporated to 20% (water phase). The samples were hydrolyzed by addition of 25 μl 5M HCl and incubation for 1 hour at 90°C. Hydrolyzed samples were neutralized with 25 μl 5M NaOH and re-extracted with 500 μl ethyl acetate. The organic phase was transferred to a new tube and 50 μl 0.5% formic acid was added. The sample was concentrated by evaporation to an approximate volume of 50 μl, after which 50 μl methanol with 0.5% formic acid was added. Before LC/MS analysis, samples were spun to remove any debris. For absolute quantitation of SA with a calibration/response-curve, SA was spiked into unrelated Col-0 leaf samples before extraction in a concentration series. Samples were separated by UHPLC on a Poroshell 120 column (2.1 x 100 mm, 1.9 micron pore) with pentafluorophenyl chemistry. Mobile phases were: A) 5% methanol, 0.5% formic acid, 10 mM ammonium formate in MilliQ; B) 0.5% formic acid in acetonitrile. Separation sequence was 20% B for 2.5’, increasing to 50% B at 5’ and increasing to 95% B at 6’. Eluted metabolites were analyzed on an Agilent 6530 Q-TOF LC/MS in negative ion mode, with dual EJS/ESI ionization. Quantitation was performed by quantifying peak areas from extracted ion chromatograms of unfragmented parent ions (for SA m/z 137.023). Peak areas were corrected with internal standard and sample fresh weight, and converted to absolute amounts using the external SA spike-in calibration curve

### Growth measurements

Rosette size was determined from topview images captured in a FluorCam 1300 system with Fluorcam v10 software (Photo Systems Instruments, Czech Republic).

### Measurement of the reactive oxygen species (ROS) burst

ROS burst assays were performed as in Johanndrees, et al. ^42^.

### qPCR analysis of gene expression

Leaves of 4.5-week-old plants were syringe-infiltrated with mock (10 mM MgCl2, 0.01% DMSO), 1 μM flg22, or 1 μM nlp24. After 1 h of treatment, leaves were harvested, snap-frozen in liquid nitrogen, and stored at -80°C. Tissue was homogenized in a TissueLyserII using 3mm glass beads. We used Spectrum Plant Total RNA kit (Sigma Aldrich) to extract total RNA. cDNA was synthesized with RevertAid H-minus reverse transcriptase (ThermoFisher) and oligo(dT)16 according to manufacturer’s instructions. qRT-PCR was performed with iTaq Universal SYBR Green Supermix (Bio-Rad) according to manufacturer’s instructions on a ViiA 7 system (ThermoFisher). Primers are listed in **Supplementary Table 1**. qRT-PCR analysis was performed by averaging Ct values of technical replicates and calculating ΔCt per sample by subtracting Ct of target gene from the Ct of the *ACTIN2* gene (AT3G18780). Statistics and visualization was based on these ΔCt values.

### Data analysis and visualization

All data analyses were performed in Spyder IE for Python v3 using *pandas* 1.4.1 package for data handling and *seaborn* 0.11.2 and *matplotlib* 3.5.1 packages for data visualization. Datasets were verified to have a normal distribution with Shapiro-Wilk test of normality (*p* > 0.05), but were allowed to have unequal variances (Bartlett’s test). Shapiro-Wilk, Bartlett’s tests, Student’s t-tests, Pearson’s correlation analysis were performed with *SciPy* 1.7.1. ANOVA, Tukey’s Post-Hoc analyses, and FDR (Benjamini-Hochberg) multiple test correction were performed with *statsmodels* 0.12.2. Significance groups were determined based on Piepho ^43^ implemented in a Python script.

## Notes

### Competing Interest Statement

The authors have declared no competing interest.

## References

1 Pieterse, C. M., Leon-Reyes, A., Van der Ent, S. & Van Wees, S. C. Networking by small-molecule hormones in plant immunity. Nat Chem Biol 5, 308–316 (2009). https://doi.org:10.1038/nchembio.164

2 Ding, P. & Ding, Y. Stories of Salicylic Acid: A Plant Defense Hormone. Trends Plant Sci 25, 549–565 (2020). https://doi.org:10.1016/j.tplants.2020.01.004

3 Hickman, R. et al. Transcriptional Dynamics of the Salicylic Acid Response and its Interplay with the Jasmonic Acid Pathway. bioRxiv, 742742 (2019). https://doi.org:10.1101/742742

4 Huot, B., Yao, J., Montgomery, B. L. & He, S. Growth–Defense Tradeoffs in Plants: A Balancing Act to Optimize Fitness. Mol Plant 7, 1267–1287 (2014). https://doi.org:10.1093/mp/ssu049

5 van Butselaar, T. & Van den Ackerveken, G. Salicylic Acid Steers the Growth-Immunity Tradeoff. Trends Plant Sci 25, 566–576 (2020). https://doi.org:10.1016/j.tplants.2020.02.002

6 Zeilmaker, T. et al. DOWNY MILDEW RESISTANT 6 and DMR6-LIKE OXYGENASE 1 are partially redundant but distinct suppressors of immunity in Arabidopsis. The Plant Journal 81, 210–222 (2015). https://doi.org:10.1111/tpj.12719

7 Zhang, K., Halitschke, R., Yin, C., Liu, C. J. & Gan, S. S. Salicylic acid 3-hydroxylase regulates Arabidopsis leaf longevity by mediating salicylic acid catabolism. Proc Natl Acad Sci 110, 14807–14812 (2013). https://doi.org:10.1073/pnas.1302702110

8 Zhang, Y. et al. S5H/DMR6 Encodes a Salicylic Acid 5-Hydroxylase That Fine-Tunes Salicylic Acid Homeostasis. Plant Physiol 175, 1082–1093 (2017). https://doi.org:10.1104/pp.17.00695

9 Bartsch, M. et al. Accumulation of isochorismate-derived 2,3-dihydroxybenzoic 3-O-beta-D-xyloside in arabidopsis resistance to pathogens and ageing of leaves. J Biol Chem 285, 25654–25665 (2010). https://doi.org:10.1074/jbc.M109.092569

10 Tripathi, J. N., Ntui, V. O., Shah, T. & Tripathi, L. CRISPR/Cas9-mediated editing of DMR6 orthologue in banana (Musa spp.) confers enhanced resistance to bacterial disease. Plant Biotechnol J 19, 1291–1293 (2021). https://doi.org:10.1111/pbi.13614

11 Kieu, N. P., Lenman, M., Wang, E. S., Petersen, B. L. & Andreasson, E. Mutations introduced in susceptibility genes through CRISPR/Cas9 genome editing confer increased late blight resistance in potatoes. Sci Rep 11, 4487 (2021). https://doi.org:10.1038/s41598-021-83972-w

12 Thomazella, D. P. T. et al. Loss of function of a DMR6 ortholog in tomato confers broad-spectrum disease resistance. Proc Natl Acad Sci 118, e2026152118 (2021). https://doi.org:10.1073/pnas.2026152118

13 Pirrello, C. et al. Grapevine DMR6-1 Is a Candidate Gene for Susceptibility to Downy Mildew. Biomolecules 12 (2022). https://doi.org:10.3390/biom12020182

14 Parajuli, S. et al. Editing the CsDMR6 gene in citrus results in resistance to the bacterial disease citrus canker. Hortic Res 9, uhac082 (2022). https://doi.org:10.1093/hr/uhac082

15 Low, Y. C., Lawton, M. A. & Di, R. Validation of barley 2OGO gene as a functional orthologue of Arabidopsis DMR6 gene in Fusarium head blight susceptibility. Sci Rep 10, 9935 (2020). https://doi.org:10.1038/s41598-020-67006-5

16 Liang, B. et al. Salicylic Acid Is Required for Broad-Spectrum Disease Resistance in Rice. Int J Mol Sci 3 (2022). https://doi.org:10.3390/ijms23031354

17 Hasley, J. A. R., Navet, N. & Tian, M. CRISPR/Cas9-mediated mutagenesis of sweet basil candidate susceptibility gene ObDMR6 enhances downy mildew resistance. PLoS One 16, e0253245 (2021). https://doi.org:10.1371/journal.pone.0253245

18 Cai, Z. et al. Generation of the salicylic acid deficient Arabidopsis via a synthetic salicylic acid hydroxylase expression cassette. Plant Methods 18, 89 (2022). https://doi.org:10.1186/s13007-022-00922-x

19 van Damme, M., Huibers, R. P., Elberse, J. & Van den Ackerveken, G. Arabidopsis DMR6 encodes a putative 2OG-Fe(II) oxygenase that is defense-associated but required for susceptibility to downy mildew. The Plant Journal 54, 785–793 (2008). https://doi.org:10.1111/j.1365-313X.2008.03427.x

20 Furniss, J. J. et al. Proteasome-associated HECT-type ubiquitin ligase activity is required for plant immunity. PLoS Pathog 14, e1007447 (2018). https://doi.org:10.1371/journal.ppat.1007447

21 Ding, Y. et al. Opposite Roles of Salicylic Acid Receptors NPR1 and NPR3/NPR4 in Transcriptional Regulation of Plant Immunity. Cell 173, 1454–1467 (2018). https://doi.org:10.1016/j.cell.2018.03.044

22 Van Wees, S. C., Van Pelt, J. A., Bakker, P. A. & Pieterse, C. M. J. Bioassays for assessing jasmonate-dependent defenses triggered by pathogens, herbivorous insects, or beneficial rhizobacteria. Methods Mol Biol 1011, 35–49 (2013). https://doi.org:10.1007/978-1-62703-414-2_4

23 Bjornson, M., Kajala, K., Zipfel, C. & Ding, P. Low-cost and High-throughput RNA-seq Library Preparation for Illumina Sequencing from Plant Tissue. Bio Protoc 10, e3799 (2020). https://doi.org:10.21769/BioProtoc.3799

24 Ewels, P., Magnusson, M., Lundin, S. & Kaller, M. MultiQC: summarize analysis results for multiple tools and samples in a single report. Bioinformatics 32, 3047–3048 (2016). https://doi.org:10.1093/bioinformatics/btw354

25 Bolger, A. M., Lohse, M. & Usadel, B. Trimmomatic: a flexible trimmer for Illumina sequence data. Bioinformatics 30, 2114–2120 (2014). https://doi.org:10.1093/bioinformatics/btu170

26 Bray, N. L., Pimentel, H., Melsted, P. & Pachter, L. Near-optimal probabilistic RNA-seq quantification. Nat Biotechnol 34, 525–527 (2016). https://doi.org:10.1038/nbt.3519

27 Soneson, C., Love, M. I. & Robinson, M. D. Differential analyses for RNA-seq: transcript-level estimates improve gene-level inferences. F1000Research 4, 1521 (2015). https://doi.org:10.12688/f1000research.7563.1

28 Love, M. I., Huber, W. & Anders, S. Moderated estimation of fold change and dispersion for RNA-seq data with DESeq2. Genome Biol 15, 550 (2014). https://doi.org:10.1186/s13059-014-0550-8

29 Lex, A., Gehlenborg, N., Strobelt, H., Vuillemot, R. & Pfister, H. UpSet: Visualization of Intersecting Sets. IEEE Trans Vis Comput Graph 20, 1983–1992 (2014). https://doi.org:10.1109/TVCG.2014.2346248

30 Bindea, G. et al. ClueGO: a Cytoscape plug-in to decipher functionally grouped gene ontology and pathway annotation networks. Bioinformatics 25, 1091–1093 (2009). https://doi.org:10.1093/bioinformatics/btp101

31 McLeay, R. C. & Bailey, T. L. Motif Enrichment Analysis: a unified framework and an evaluation on ChIP data. BMC Bioinformatics 11, 165 (2010). https://doi.org:10.1186/1471-2105-11-165

32 O’Malley, R. C. et al. Cistrome and Epicistrome Features Shape the Regulatory DNA Landscape. Cell 165, 1280–1292 (2016). https://doi.org:10.1016/j.cell.2016.04.038

33 Franco-Zorrilla, J. M. et al. DNA-binding specificities of plant transcription factors and their potential to define target genes. Proc Natl Acad Sci 111, 2367–2372 (2014). https://doi.org:10.1073/pnas.1316278111

34 Katoh, K., Misawa, K., Kuma, K. & Miyata, T. MAFFT: a novel method for rapid multiple sequence alignment based on fast Fourier transform. Nucleic Acids Res 30, 3059–3066 (2002). https://doi.org:10.1093/nar/gkf436

35 Minh, B. Q. et al. IQ-TREE 2: New Models and Efficient Methods for Phylogenetic Inference in the Genomic Era. Mol Biol Evol 37, 1530–1534 (2020). https://doi.org:10.1093/molbev/msaa015

36 Kalyaanamoorthy, S., Minh, B. Q., Wong, T. K. F., von Haeseler, A. & Jermiin, L. S. ModelFinder: fast model selection for accurate phylogenetic estimates. Nat Methods 14, 587–589 (2017). https://doi.org:10.1038/nmeth.4285

37 Shiu, S. H. et al. Comparative analysis of the receptor-like kinase family in Arabidopsis and rice. Plant Cell 16, 1220–1234 (2004). https://doi.org:10.1105/tpc.020834

38 Restrepo-Montoya, D., Brueggeman, R., McClean, P. E. & Osorno, J. M. Computational identification of receptor-like kinases “RLK” and receptor-like proteins “RLP” in legumes. BMC Genomics 21, 459 (2020). https://doi.org:10.1186/s12864-020-06844-z

39 Wang, G. et al. A genome-wide functional investigation into the roles of receptor-like proteins in Arabidopsis. Plant Physiol 147, 503–517 (2008). https://doi.org:10.1104/pp.108.119487

40 Li, P. et al. RGAugury: a pipeline for genome-wide prediction of resistance gene analogs (RGAs) in plants. BMC Genomics 17, 852 (2016). https://doi.org:10.1186/s12864-016-3197-x

41 Letunic, I. & Bork, P. Interactive Tree Of Life (iTOL) v5: an online tool for phylogenetic tree display and annotation. Nucleic Acids Res 49, W293–W296 (2021). https://doi.org:10.1093/nar/gkab301

42 Johanndrees, O. et al. Variation in plant Toll/Interleukin-1 receptor domain protein dependence on ENHANCED DISEASE SUSCEPTIBILITY 1. Plant Physiol, kiac480 (2022). https://doi.org:10.1093/plphys/kiac480

43 Piepho, H.-P. An Algorithm for a Letter-Based Representation of All-Pairwise Comparisons. Journal of Computational and Graphical Statistics 13, 456–466 (2004). https://doi.org:10.1198/1061860043515

