## Supplemental Figures and Tables for "Basal expression of immune receptor genes requires low levels of the phytohormone salicylic acid"

### Supplementary Figures

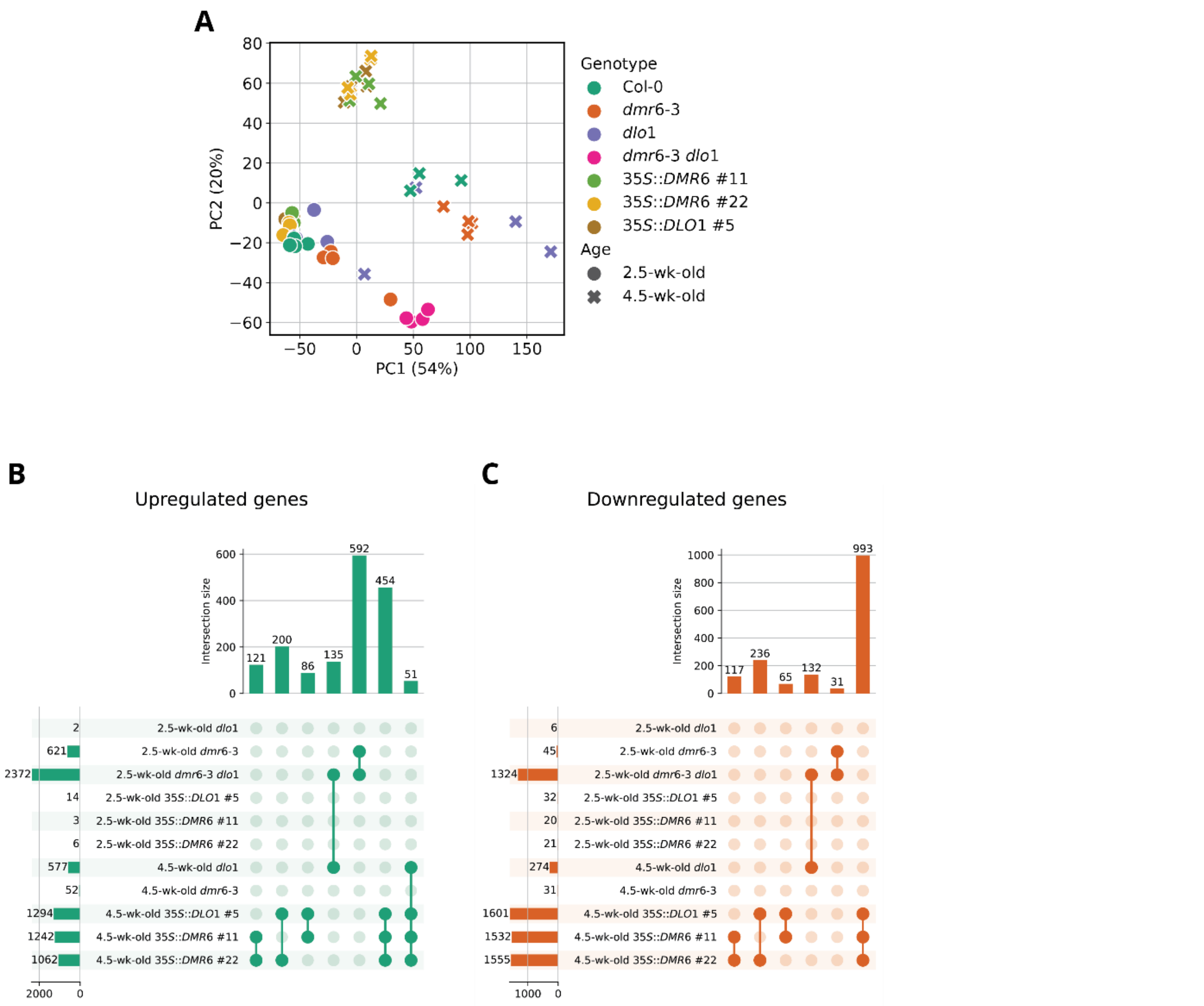


**Supplementary Figure 1. Transcriptome analysis of plants with perturbed SA catabolism. A)** Principal component analysis (PCA) plot of normalized transcript counts of 19,405 expressed genes. The percentage explained by each principal component (PC) is indicated. PC1 mainly separates different genotypes, while PC2 mainly separates plant age. **B)** Upset plots of genes significantly ($|\log_2 FC| \geq 1$, FDR-adj. $p \leq 0.05$) up (left) or down (right) regulated in each genotype compared to wild-type. Indicated in the bar graphs on the left are the number of DEG in that genotype. Connected points are groups of DEG that are regulated similarly in connected genotypes. The size of these groups (*intersection size*) are displayed in the bar graph at the top. Gene groups smaller than 25 were excluded.

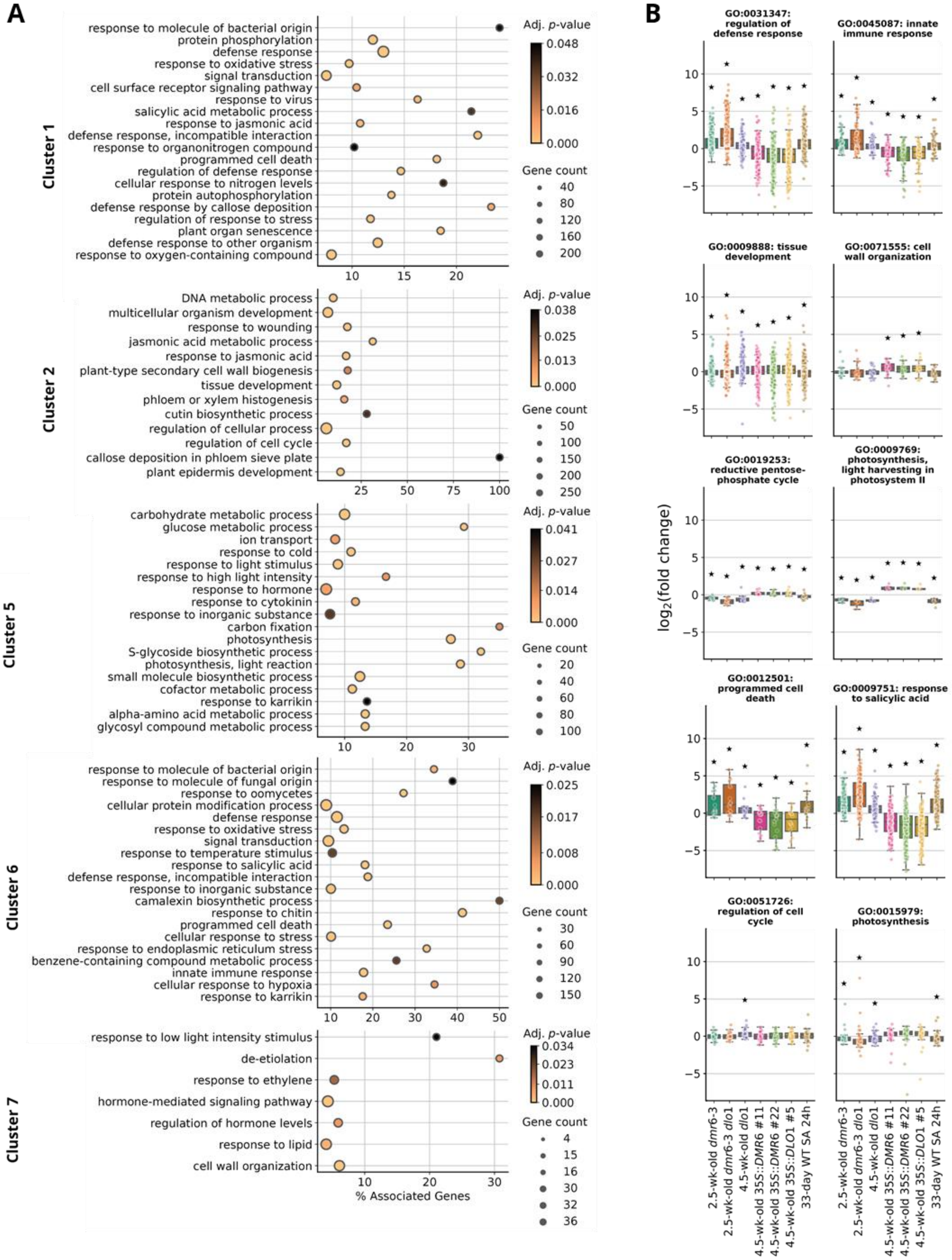


**Supplementary Figure 2. GO term enrichment analysis of gene clusters in Figure 1D. A)** Dotplots of GO terms significantly enriched in clusters with more than 500 genes. The color of the dots indicates the *p*-value from the enrichment test, size of the dots indicates the number of genes in the cluster that belong to that GO term. Position on the *x*-axis indicates relative enrichment as the percentage of genes in the cluster over the total number of genes belonging to that GO term. **B)** Boxplots with $\log_2$ fold change values of all expressed genes belonging to several immunity or growth and development-related GO terms. Asterisks denote expression with significant deviation from a hypothetical mean of zero, Student's one sample t-test + FDR (Benjamini-Hochberg) $p \leq 0.05$.

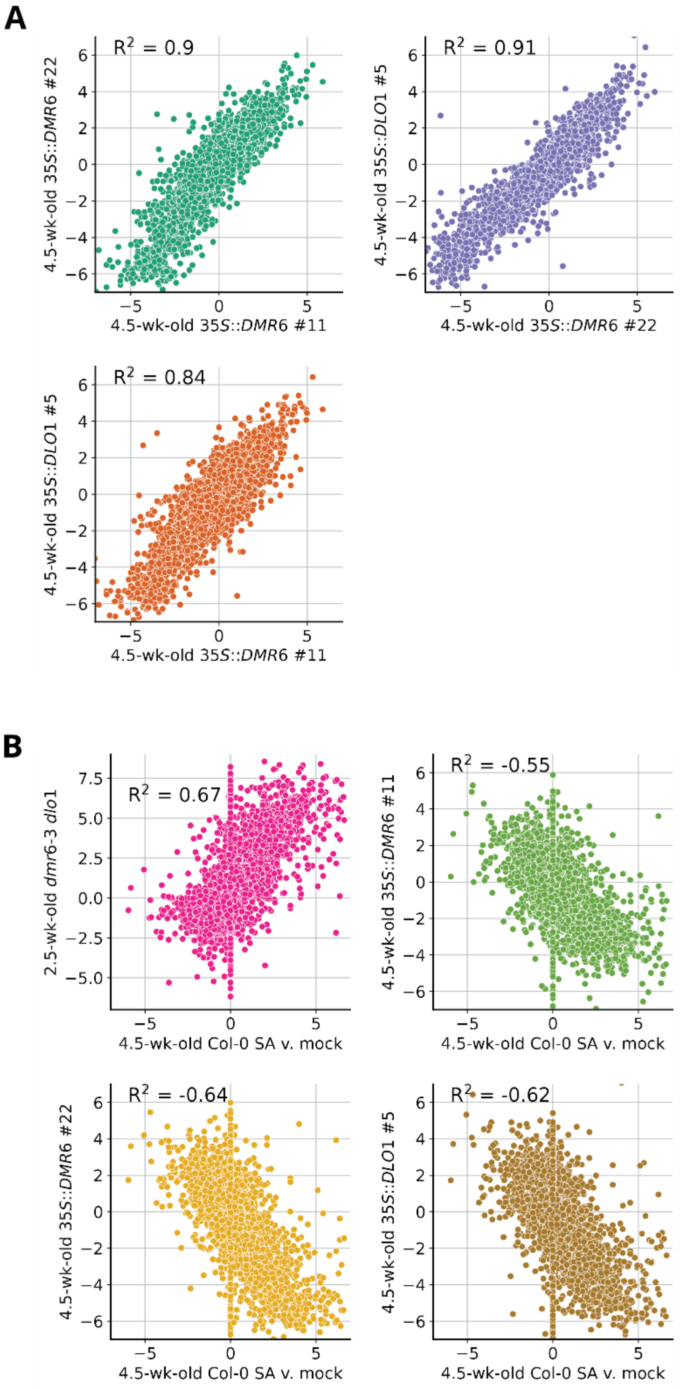


**Supplementary Figure 3. Correlation analysis** between **(A)** SA-treated Col-0 and *DMR6/DLO1* mutants and overexpression lines, and **(B)** 4.5-week-old *DMR6/DLO1* overexpression lines. Depicted are $\log_2|FC|$ values for all 6234 DEG ($|\log_2 FC| \geq 1$, FDR-adj. $p \leq 0.05$). Figures are annotated with Pearson's correlation metrics. $p$-values for Pearson correlation distances were < 0.001

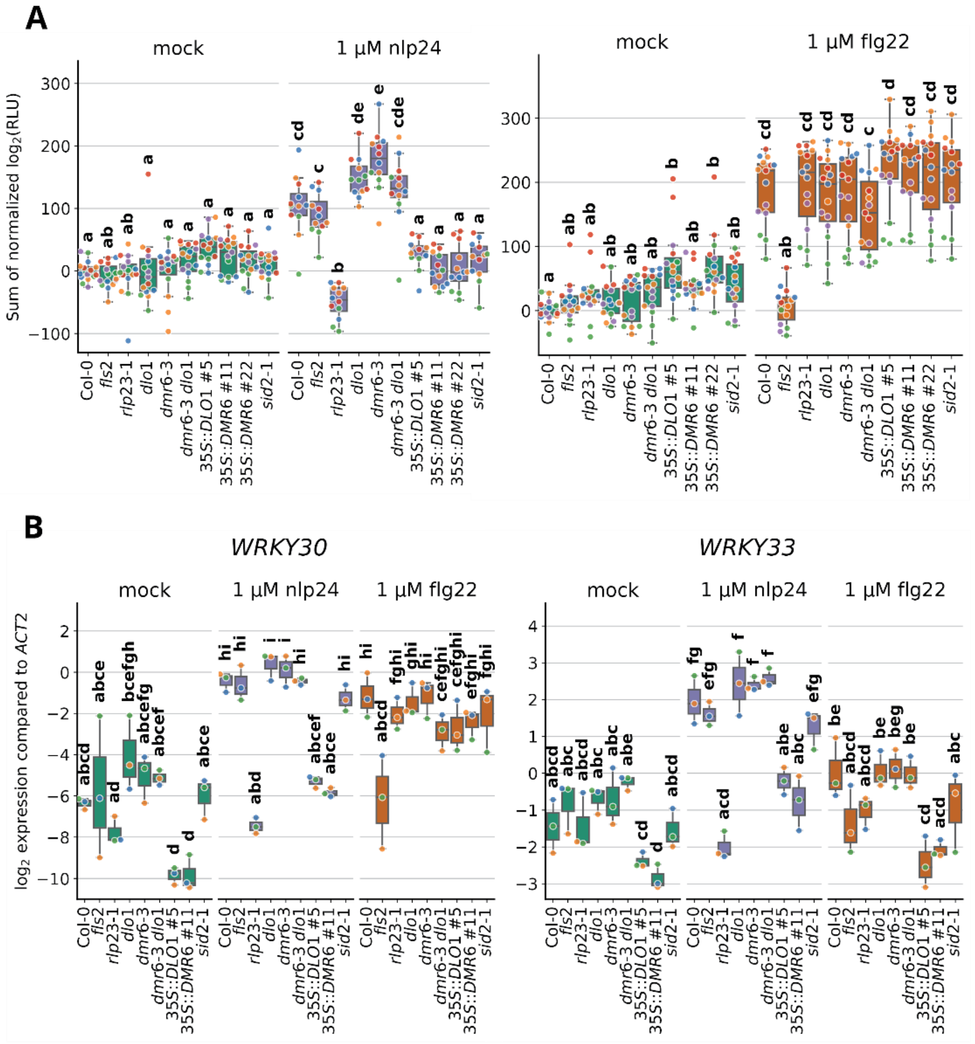


**Supplementary Figure 4. PTI responses in *dmr6-3/dlo1* mutants and *DMR6/DLO1* overexpression lines. A)** ROS burst responses to nlp24 were reduced in SA-depleted genotypes, but enhanced only in *dmr6-3* single mutant plants. RLU: relative light units, as the sum of area under the curve over 86 minutes after treatment. **B)** *WRKY30* and *WRKY33* expression changes were reduced in *DMR6/DLO1* overexpression lines after the nlp24 treatment. Transcript levels were measured by qRT-PCR at three hours after the inoculation with PAMP or mock. **A and B)** Data was derived from two independent experiments, as indicated by differently colored dots. Different letters denote statistically significant differences between the groups, from two-way ANOVA + Tukey's Post-Hoc test, $p \leq 0.05$.

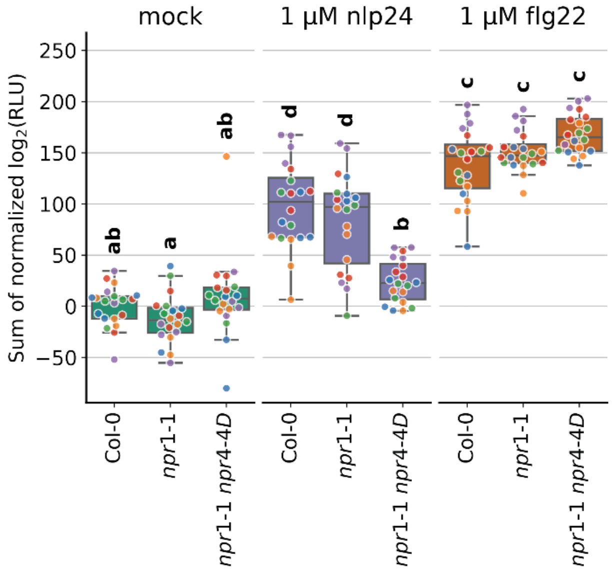


**Supplementary Figure 5. Attenuation of PTI responses in SA-insensitive mutants.** ROS burst responses to nlp24 are reduced in *npr1-1 npr4-4D* plants. RLU: relative light units, as sum of area under the curve over 61 minutes after treatment. The plants were 4.5 weeks old. Data are derived from three independent experiments, as indicated by differently colored dots. Different letters denote statistically significant differences between groups, from two-way ANOVA + Tukey's Post—Hoc, $p \leq 0.05$.

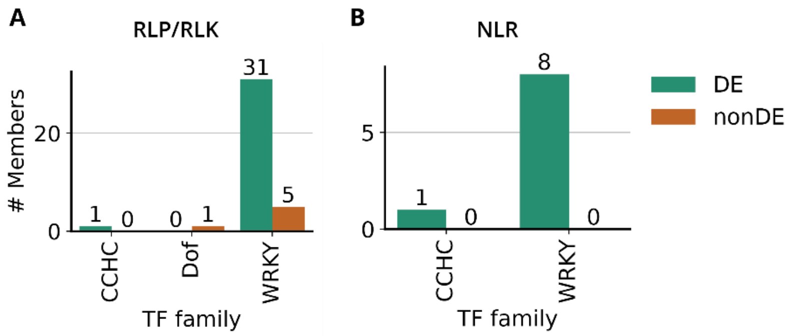


**Supplementary Figure 6.** **Enrichment of WRKY TF binding sites in SA-responsive *RLK/RLP* or *NLR* gene promoters.** Barplot describing the number of TF family members with predicted enrichment of TF binding sites in promoters of *RLP/RLKs* **(A)** or all *NLRs*[22] **(B)** that were downregulated in 4.5-week-old *DMR6/DLO1* overexpression lines and upregulated in the 2.5-week-old *dmr6-3 dlo1* double mutant. *RLP/RLKs* or *NLRs* that were not differentially expressed in these genotypes are marked as 'nonDE'.

**Supplementary Table 1. General information about the RNAseq samples. A)** Samples generated in this study with descriptive metadata. *reads_processed* and *reads_pseudoaligned* are the number of reads that passed quality filtering by *trimmomatic* and were successfully pseudoaligned by *kallisto*, respectively. **B)** Normalized pseudocounts output from *deseq2* analysis for all 19405 expressed genes per sample.

**Supplementary Table 2. Overview of differentially expressed genes. A)** Differential expression calling for all 19405 expressed genes for each genotype to its respective wild-type control. 1: significantly upregulated, -1: significantly downregulated, 0: not significantly differentially expressed. **B)** Normalized log2|FC| values for all expressed genes from *deseq2* output. **C)** Assignment of cluster IDs for 6234 DEGs in this study.

**Supplementary Table 3. GO term enrichment analysis for each cluster.** Output from ClueGO on GO terms of type Biological Process enriched in the identified clusters.

**Supplementary Table 1. Primers used in this study.**

| Name | Gene code | Sequence 5' → 3' | Use |
|---|---|---|---|
| **nDL041_WRKY30_Fw** | AT5G24110 | AGCCAAATTTCCAAGAGGAT | qRT-PCR |
| **nDL042_WRKY30_Rv** | AT5G24110 | GCAGCTTGAGAGCAAGAATG | qRT-PCR |
| **nDL079_WRKY33_Fw** | AT2G38470 | GGCTCATCGATTGTCAGCAG | qRT-PCR |
| **nDL079_WRKY33_Rv** | AT2G38470 | CCTTAACGACTTTCTGGCCG | qRT-PCR |
| **oTvB496_qACT2_Fw** | AT3G18780 | AATCACAGCACTTGCACCA | qRT-PCR |
| **oTvB497_qACT2_Rv** | AT3G18780 | GAGGGAAGCAAGAATGGAAC | qRT-PCR |
| **EnrichS1** | | AATGATACGGCGACCACCGA | Enrichment primer |
| **EnrichS2** | | CAAGCAGAAGACGGCATACGA | Enrichment primer |
| **RNAseq-D501** | | AATGATACGGCGACCACCGAGATCTACACTATAGCCTACACTCTTTCCCTACACGACGCTCTTCCGATCT | i5 Indexed primer |
| **RNAseq-D502** | | AATGATACGGCGACCACCGAGATCTACACATAGAGGCACACTCTTTCCCTACACGACGCTCTTCCGATCT | i5 Indexed primer |
| **RNAseq-D503** | | AATGATACGGCGACCACCGAGATCTACACCCTATCCTACACTCTTTCCCTACACGACGCTCTTCCGATCT | i5 Indexed primer |
| **RNAseq-D504** | | AATGATACGGCGACCACCGAGATCTACACGGCTCTGAACACTCTTTCCCTACACGACGCTCTTCCGATCT | i5 Indexed primer |
| **RNAseq-D505** | | AATGATACGGCGACCACCGAGATCTACACAGGCGAAGACACTCTTTCCCTACACGACGCTCTTCCGATCT | i5 Indexed primer |
| **RNAseq-D506** | | AATGATACGGCGACCACCGAGATCTACACTAATCTTAACACTCTTTCCCTACACGACGCTCTTCCGATCT | i5 Indexed primer |
| **RNAseq-D507** | | AATGATACGGCGACCACCGAGATCTACACCAGGACGTACACTCTTTCCCTACACGACGCTCTTCCGATCT | i5 Indexed primer |
| **RNAseq-D508** | | AATGATACGGCGACCACCGAGATCTACACGTACTGACACACTCTTTCCCTACACGACGCTCTTCCGATCT | i5 Indexed primer |
| **RNAseq-D701** | | CAAGCAGAAGACGGCATACGAGATATTACTCGGTGACTGGAGTTCAGACGTGTGCTCTTCCGATC | i7 Indexed primer |
| **RNAseq-D702** | | CAAGCAGAAGACGGCATACGAGATTCCGGAGAGTGACTGGAGTTCAGACGTGTGCTCTTCCGATC | i7 Indexed primer |

| | | |
|---|---|---|
| **RNAseq-D703** | CAAGCAGAAGACGGCATACGAGATCGCTCATTGTGACTGGAGTTCAGACGTGTGCTCTTCCGATC | i7 Indexed primer |
| **RNAseq-D704** | CAAGCAGAAGACGGCATACGAGATGAGATTCCGTGACTGGAGTTCAGACGTGTGCTCTTCCGATC | i7 Indexed primer |
| **RNAseq-D705** | CAAGCAGAAGACGGCATACGAGATATTCAGAAGTGACTGGAGTTCAGACGTGTGCTCTTCCGATC | i7 Indexed primer |
| **RNAseq-D706** | CAAGCAGAAGACGGCATACGAGATGAATTCGTGTGACTGGAGTTCAGACGTGTGCTCTTCCGATC | i7 Indexed primer |
| **RNAseq-D707** | CAAGCAGAAGACGGCATACGAGATCTGAAGCTGTGACTGGAGTTCAGACGTGTGCTCTTCCGATC | i7 Indexed primer |
| **RNAseq-D708** | CAAGCAGAAGACGGCATACGAGATTAATGCGCGTGACTGGAGTTCAGACGTGTGCTCTTCCGATC | i7 Indexed primer |
| **RNAseq-D709** | CAAGCAGAAGACGGCATACGAGATCGGCTATGGTGACTGGAGTTCAGACGTGTGCTCTTCCGATC | i7 Indexed primer |
| **RNAseq-D710** | CAAGCAGAAGACGGCATACGAGATTCCGCGAAGTGACTGGAGTTCAGACGTGTGCTCTTCCGATC | i7 Indexed primer |
| **RNAseq-D711** | CAAGCAGAAGACGGCATACGAGATTCTCGCGCGTGACTGGAGTTCAGACGTGTGCTCTTCCGATC | i7 Indexed primer |
| **RNAseq-D712** | CAAGCAGAAGACGGCATACGAGATAGCGATAGGTGACTGGAGTTCAGACGTGTGCTCTTCCGATC | i7 Indexed primer |
